# Phylo-Movies: Animating Phylogenetic Trees from Sliding-Window Analyses

**DOI:** 10.64898/2026.04.01.715821

**Authors:** Enes Berk Sakalli, Simon E. Haendeler, Arndt von Haeseler, Heiko A. Schmidt

## Abstract

Sliding-window phylogenetic analyses of multiple sequence alignments (MSAs) generate sequences of phylogenetic trees that can reveal recombination and other sources of phylogenetic conflict, yet comparing trees across genomic windows remains challenging. Phylo-Movies is a browser-based tool, also available as a standalone desktop application, that decomposes topological differences between consecutive phylogenetic trees into interpretable subtree migrations and animates these transformations. We demonstrate its utility in two contexts: identifying recombintion breakpoints in norovirus genomes, where lineages shift from polymerase-based to capsid-based clustering at the ORF1/ORF2 junction, and detecting rogue taxa that change position across bootstrap replicates. Phylo-Movies complements summary statistics such as Robinson–Foulds distances by showing which lineages move, where they move from, and which new groupings they form. Phylo-Movies is freely available at https://github.com/enesberksakalli/phylo-movies, with a norovirus demonstration video at https://vimeo.com/1162400544, the first rogue taxon example at https://vimeo.com/1162561152, and the second example at https://vimeo.com/1162563101.

## 1 Introduction

Ideally, phylogenetic inference of whole-genome analysis reveals that the entire genomes evolved under a single evolutionary history. Reality, however, is more complex. For example, if two related viruses infect a single cell, during genome amplification RNA-dependent RNA polymerase (RdRp) can change the template from one viral genome to the other, commonly referred to as template switching (Simon-Lorière and Holmes, 2011). This process links parts of the co-infecting genomes, generating a hybrid virus variant. The different regions of this recombinant genome now reflect the different evolutionary histories of their origin. Similar effects occur via lateral gene transfer between bacteria or crossover during meiosis in eukaryotes (Ochman *et al*., 2000; Hunter, 2015).

Recombination between highly divergent sequences may disrupt essential gene functions, so successful recombinants typically arise from homologous regions where sequence and gene order are conserved. Consequently, recombinant genomes usually retain enough similarity to be captured in a single multiple sequence alignment (MSA). However, treating such an alignment as a single evolutionary unit can obscure conflicting phylogenetic signals (Posada and Crandall, 2002). Sliding-window approaches address this problem by reconstructing a phylogenetic tree for each successive, overlapping window along the alignment (Salminen *et al*., 1995; Lole et al., 1999) (Fig. 1a,b). Because each window samples only a short genomic segment of the alignment, its tree reflects the local evolutionary history of that region alone. Where no recombination has occurred, consecutive windows yield similar or identical topologies (i.e., branching structures independent of branch lengths); where recombination has occurred, the topology shifts to reflect the distinct ancestry of the recombinant segment, and the genomic position at which this shift occurs approximates the location of a recombination breakpoint.

**Figure 1:**
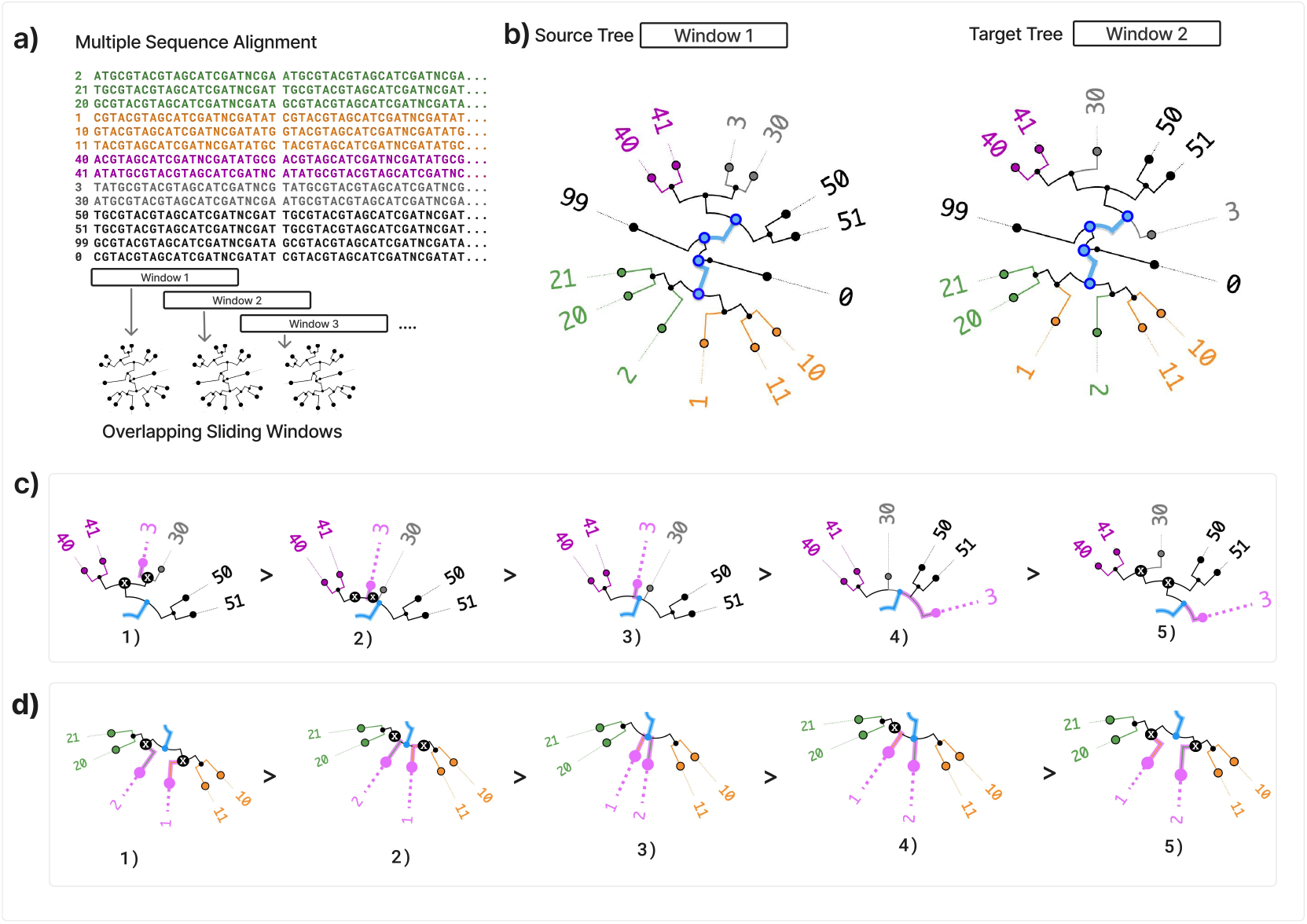
(a) A genomic alignment is divided into overlapping sliding windows of defined length and step size. For each window, a phylogenetic tree is reconstructed, generating a sequence of trees for Phylo-Movies. (b) Two consecutive trees (windows 1 and 2) are shown as source and target trees. Sequences and corresponding tree nodes are coloured according to the grouping in the source tree. Two sequential topological changes (upper and lower parts) are illustrated in (c) and (d). (c) In the upper example, leaf 3 changes position between source and target. In the source tree, leaf 3 forms a subtree with leaf 30. In the target tree, leaf 3 instead joins the subtree containing leaves 50 and 51. The transformation proceeds relative to the Pivot Edge (thick light blue). First, branches unique to the source tree collapse, producing a temporary multifurcation. Leaf 3 is then repositioned within this multifurcation to join the 50/51 subtree, after which branches unique to the target tree are inserted, yielding the target topology. (d) The lower example shows two independent rearrangements. In the source tree, leaves 1 and 2 belong to the same subtree. In the target tree, leaf 1 leaves this subtree and joins the subtree containing leaves 20 and 21, while leaf 2 is repositioned to form a new subtree. As above, source-unique branches collapse, the taxa are repositioned within the resulting multifurcation, and target-specific branches are inserted to obtain the final topology.

As the window progresses, the tree topologies change to reflect the local evolutionary history, producing a sequence of trees ordered by genomic position. Topological shifts in this sequence mark candidate recombination breakpoints. This approach requires sufficient sequence divergence: recombination between nearly identical parents leaves the tree structure unchanged and thus escapes detection.

Several tools exist to analyse recombinant alignments. RDP5, SimPlot, SimPlot++, and TOPALi focus primarily on detecting recombinants and locating breakpoints (Martin *et al*., 2020; Lole et al., 1999; Samson et al., 2022; Milne et al., 2004). Tree House Explorer takes a complementary approach, showing the distribution of tree topologies across genomic windows in a genome browser (Harris *et al*., 2022). However, it does not visualise how trees from consecutive windows relate to each other, leaving users unable to identify which subtrees rearrange, where they move, or which new groupings they form.

A phylogenetic tree represents inferred relationships among aligned sequences, with its topology describing how taxa group relative to one another. Comparing topologies from adjacent windows can therefore reveal when a lineage clusters with one group in one genomic region but with a different group in another—a signature of recombination.

The scale of modern datasets compounds this challenge. Small step sizes generate hundreds or thousands of largely redundant trees that must be inspected for subtle rearrangements, whereas larger step sizes can produce abrupt topological changes between consecutive windows, making it difficult to track how specific lineages move. For example, sliding-window analysis of a 10 kb HIV genome with a 10 nt step produces approximately 1,000 trees (Abecasis *et al*., 2007). Identifying the windows where tree rearrangements occur requires tracing branching structures across hundreds of trees, a task poorly suited to static visualisation (Graham and Kennedy, 2009).

To address these challenges, we developed Phylo-Movies, that animates topological rearrangements between consecutive phylogenetic trees. Users may supply a pre-computed tree sequence or upload an MSA, from which trees are inferred using FastTree (Price *et al*., 2010) with user-defined window and step sizes and then visualised with Phylo-Movies. Trees can be rooted using either an outgroup or midpoint rooting. Phylo-Movies visualises topological changes by representing the transition between two consecutive trees as a series of Subtree Prune and Regraft (SPR) operations. An SPR operation removes (prunes) a subtree from one location in a rooted tree and reattaches (regrafts) it to another branch, yielding a new topology after a single rearrangement (Bordewich and Semple, 2005). Each such move is animated with the affected subtree highlighted, allowing regions of substantial topological change within the tree sequence to be readily identified. We next describe the algorithmic framework underlying the animation and present two use cases illustrating its analytical value.

## 2 The Phylo-Movies Approach

### 2.1 How to Animate Topological Changes Between Trees

Phylo-Movies highlights and animates topological changes within a sequence of phylogenetic trees. Visualisation requires rooted trees, which can be obtained using either an outgroup or midpoint rooting (Edwards, 2019). Trees are displayed using a circular layout (cf. Felsenstein, 2004, chap. 36).

To visualise the whole sequence of trees, topological changes are animated between each pair of consecutive trees, transforming the source tree into the succeeding target tree (cf. Fig. 1b). To define this transformation, we use the concepts of splits and consensus trees (cf. Bryant, 1997).

Every internal branch defines a (non-trivial) split that partitions the leaves into two subsets. Comparing the source and target trees identifies three categories of splits: splits shared by both, splits unique to the source tree, and splits unique to the target tree. The Robinson–Foulds distance (RF) between the two trees is defined as the number of splits unique to the source plus the number unique to the target (Robinson and Foulds, 1981, 1979).

To visualise the transition from the source tree to the target tree, Phylo-Movies uses the strict consensus tree, which contains only the splits present in both trees, with branch lengths set to the mean of the corresponding values in the source and target trees. The transformation is organised around a **Pivot Edge**, defined as a branch shared by both trees whose immediate descendant branches differ between the source tree and the target tree.

This process is shown in the workflow in Fig. 1. From the alignment in Fig. 1a, window subalignments are extracted to reconstruct the trees in Fig. 1b. Fig. 1c shows a topological rearrangement in the upper part of the trees, namely the migration of **leaf 3** (marked magenta). In the source tree (1, Fig. 1c), leaf 3 is sister to leaf 30. To reach the target tree topology (5, Fig. 1c), where leaf 3 groups with the (30, 40,41, 50, 51) subtree, the algorithm identifies the **Pivot Edge** (thick blue arc) and performs the rearrangement relative to it.

The transformation begins by shrinking to zero the branch leading to the subtree of leaves 3 and 30, as well as the branch rooting the subtree containing 3, 30, 40, and 41 (Fig. 1c. 1). Both branches correspond to splits unique to the source tree and remain visible in tree 2) as black dots. These zero-length branches are then removed, yielding the intermediate strict consensus tree (Fig. 1c.1). Subsequently, leaf 3 moves to its correct position within the multifurcation (Fig. 1c.4), and finally the two new branches unique to the target tree are inserted and expanded in Fig. 1c.5: the branch rooting subtree (30, 40, 41) and the branch rooting subtree (30, 40, 41, 50, 51).

Fig. 1d presents a more complex scenario where two taxa (**leaf 1** and **leaf 2**, both marked magenta) are moving. In the source tree 1), leaf 1 clusters with leaves 10 and 11, while leaf 2 clusters with 20 and 21. To transform towards the target tree 5), the branches rooting the subtrees (1, 10, 11) and (2, 20, 21) are shrunk to zero length, see black nodes at the end of the magenta branches in tree 2). In the next step (tree 3), these zero-length branches are removed yielding the intermediate strict consensus in tree 3) where both leaves attach to the same multifurcation outside the Pivot edge. Again the branches are reordered to match the target topology (while the consensus topology remains static - see tree 3). Then, the new branch rooting subtree (1, 20, 21) is inflated (tree 4) and, finally, the branch for subtree (2, 10, 11) is inserted in tree 5. This demonstrates how multi-taxon rearrangements are decomposed into sequences of branch collapse and expansion.

For each pair of consecutive trees in the sequence, the preceding tree (source tree) is compared with the succeeding tree (target tree). Pivot Edges are defined as shared branches whose immediate descendant branches define splits unique to the source that are replaced by splits unique to the target. Identifying these points isolates the subtrees associated with the topological conflict. This corresponds to rooted Subtree Pruning and Regrafting (SPR) operations, in which a subtree is detached from one location and reattached elsewhere (Bordewich and Semple, 2005). In the animation, the Pivot Edge serves as the base from which these subtrees move to their new positions. At each Pivot Edge, subtrees that remain structurally identical between source tree and target tree are retained, and only those whose placement differs are animated.

Once the moving subtrees are identified, their relocation to the target position is animated while the consensus structure remains static. To maintain visual clarity, Phylo-Movies animates these moves sequentially. When a single moving subtree contains multiple internal conflicts, a heuristic decomposes the movement into smaller sequential transitions.

For each rearrangement, the visualisation animates the corresponding SPR move by smoothly transitioning from the source tree layout to the target tree layout, producing a sequence of frames (default: 30). To minimise extraneous motion, the tool optimises the leaf ordering of the target tree to match the source layout, ensuring that only subtrees involved in topological changes appear to move. Additional navigation and analysis features are described below.

### 2.2 Phylo-Movies Features and User Interface

The Phylo-Movies user interface (Fig. 2) consists of several synchronised panels. The *animation and tree viewer panel* (Fig. 2A) displays the transformation between trees, where moving subtrees are highlighted.

**Figure 2:**
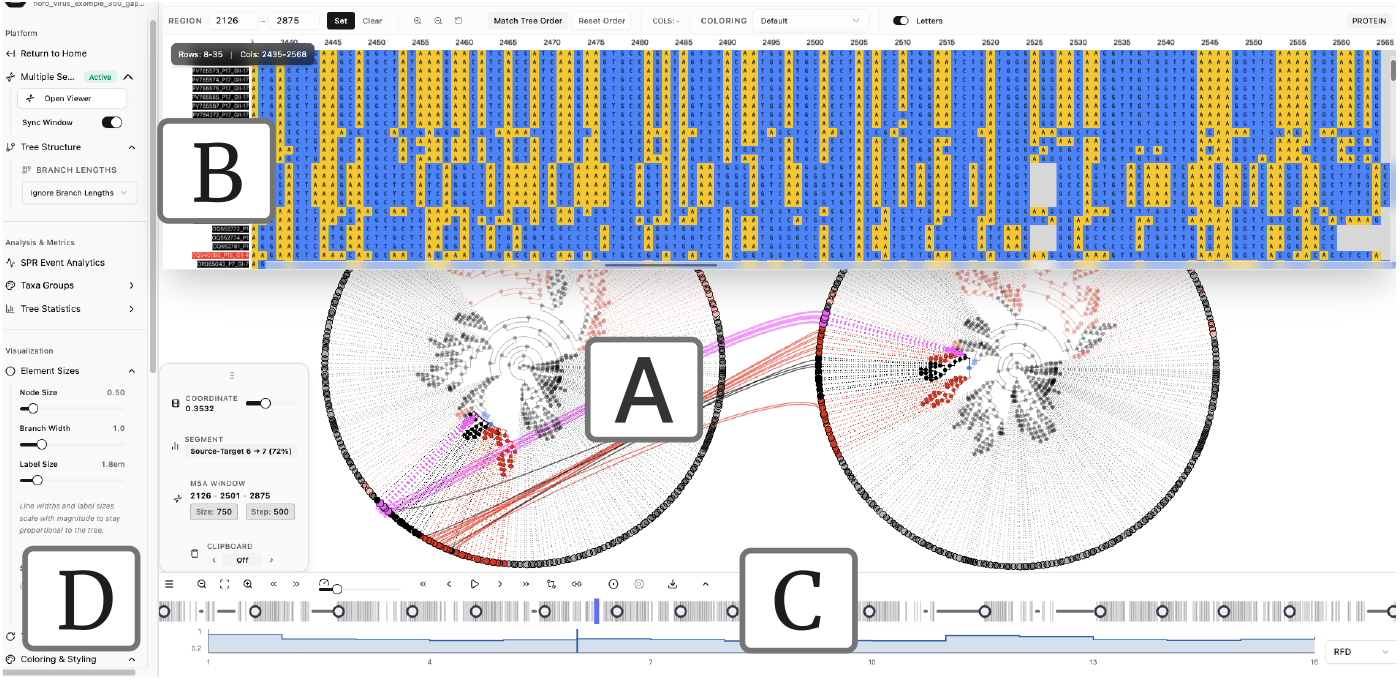
The Phylo-Movies interface for visualising phylogenetic tree sequences, showing an example analysing the Norovirus genomes. (A) **Tree viewer**: visualising the phylogeny for the current window/position. Moving subtrees are highlighted in red and cyan with their pivot edge in blue; the subtree undergoing change in the current step is coloured blue. Leaves can also be coloured by user-selected metadata-defined groups, here by polymerase genotype. Phylo-Movies can also show a reference tree, here the target tree, with selected taxa linked by lines. (B) **MSA viewer**: presents the current region of the underlying alignment, highlighting the active window, with explicit region bounds. When window synchronisation is enabled, the alignment position shown will change according to the current tree position. (C) **playback control, progress and distance panel**: The top row contains control buttons to play, stop, and step through the animation. Below you find a progress bar that shows the trees of the sequence as circles and the individual transformations as vertical bars. The vertical blue playhead can also be used to navigate through the visualisation across the genome forward or backward manually. The distance plot at the bottom helps localise interesting regions. Different measures can be chosen such as Robinson–Foulds, weighted RF or branch-length distance. The vertical bar marks the current position and can also be used to move the focus manually. (D) **Left control panel**: file/alignment actions and visualisation settings (branch-length handling, geometry/layout transforms, and taxa colouring), including the window-synchronisation toggle. **Right status & grouping panel**: progress through the current segment/window, tree scale and branch-length summaries, and group/analytics controls for colouring and exploration.

The *playback control panel* (Fig. 2C) includes buttons to play, stop, and step through the animation forward and backward. The *timeline chart* below (Fig. 2C) tracks the genomic location with a playhead. Clicking or dragging the playhead adjusts the animation position.

The distance plot below the timeline shows Robinson–Foulds distances (RF; Robinson and Foulds, 1981, 1979) between adjacent trees. Standard RF treats each internal branch as a split (a bipartition of the taxa) and counts how many splits are present in one tree but not in the other; branch lengths are ignored.

The weighted RF distance is then the sum, over all splits that appear in either tree, of the absolute difference between the two split lengths (using 0 for splits that are absent in one of the trees). Users can switch between RF, weighted RF, and branch score distance.

The *alignment panel* (Fig. 2B) displays the genomic alignment synchronised with the current tree, allowing users to associate genome positions with topological changes. A *setup panel* on the left (Fig. 2D) provides configuration options for the visualisation and toggles synchronisation between the alignment viewer and the tree state. A *control panel* on the right enables colouring of subtrees and taxa by metadata annotations and metadata-defined groups.

Phylo-Movies also supports side-by-side tree comparison: users can pin any tree from the sequence as a reference (Fig. 2A, right), helping to track where subtrees were positioned before and how they have moved relative to a chosen starting point. A comparative mode links corresponding taxa between phylogenies to highlight structural differences. Rendering is implemented via WebGL to support large trees and alignments at sufficient speed.

The visualisation supports several customisation options designed for scientific interpretability. Subtree colouring propagates a leaf colour to internal branches only when all descendants share that colour, making monophyletic groups visually distinct. Users can load metadata from a CSV file to define metadata-defined groups (e.g., genotype, host, or geographic origin) and interactively switch between grouping schemes. Node size can be scaled, and highlighted elements—pivot edges, marked subtrees, or structures from previous frames—are rendered with increased radius or thicker outlines to improve perceptual salience. Edge highlighting employs explicit visual precedence, ensuring that marked subtrees and active change edges remain distinguishable from baseline colouring.

## 3 Use Cases and Results

### 3.1 Recombination-Induced Topological Shifts in Norovirus GII

Recombination drives norovirus evolution by enabling viruses to acquire novel capsid variants while retaining a functional polymerase, thereby evading host immunity (Chhabra *et al*., 2019; Lindesmith *et al*., 2008; Bull et al., 2005). Because the polymerase gene (ORF1) and capsid gene (ORF2) frequently have distinct evolutionary origins, a dual-genotype nomenclature of the form GII.*y* [P*x*] is used, where GII.*y* denotes the capsid genotype and P*x* the polymerase genotype (Chhabra *et al*., 2019). Matching genotypes (e.g., GII.4[P4]) suggest a shared evolutionary history, whereas mismatched combinations (e.g., GII.4[P16]) indicate recombinant origins.

To investigate how tree topology shifts across the recombination breakpoint, we used Phylo-Movies to visualise a sliding-window phylogenetic analysis on a global dataset of 334 full-genome Norovirus sequences (8,058 bp alignment) from 32 countries, between the years of 1968–2025. Sequences were downloaded from the Nextstrain-curated GenBank dataset (Hadfield *et al*., 2018), subsampled using Augur (Huddleston *et al*., 2021), and genotyped using Nextclade (Aksamentov *et al*., 2021). The dataset encompasses 30 capsid and 28 polymerase genotypes, with 167 sequences (50.0%) representing recombinant strains in which the capsid and polymerase genotypes differ.

The sequences were aligned using MAFFT v7.526 (Katoh and Standley, 2013) and trimmed using trimAl (Capella-Gutiérrez *et al*., 2009). Trees were inferred using FastTree (Price *et al*., 2010) under the GTR model, with a window size of 750 bp and a step size of 500 bp, and were subsequently visualised with Phylo-Movies. All trees were rooted at the midpoint, and the Robinson–Foulds distances between adjacent windows were used to identify regions of pronounced topological change. (Fig. 2).

At the ORF1/ORF2 junction (approximately position 5,100 bp), both the Robinson–Foulds distance and the weighted Robinson–Foulds distance show pronounced peaks, indicating a major topological change accompanied by shifts in branch lengths. In ORF1 windows, sequences group by polymerase genotype; as the window progresses into ORF2, they instead group by capsid genotype.

GII.4[P16] recombinants, for instance, cluster with other P16 sequences in ORF1 but group with GII.4 sequences in ORF2, illustrating their mosaic origin: segments derived from different parental lineages form contiguous blocks that associate with distinct clusters depending on genomic position. Because the animation decomposes topological transitions into sequential SPR moves, the relocation of these subtrees can be followed step by step. We also observed a small GII.2[P16] subtree, comprising approximately 20 taxa, shifting from the P16-associated cluster in ORF1 to join other GII.2 sequences in ORF2—a rearrangement that would be difficult to detect in a static tree comparison. Similarly, GII.6[P7] variants, predominantly sampled in Asia, repeatedly grouped with South American GII.6 sequences across multiple windows, illustrating how geographic structure can be traced through the animation. This approach can also be applied to sequences of unknown origin: by including them in the analysis, one can observe which clusters they join in different genomic regions, thereby suggesting their polymerase and capsid genotype assignments.

A feature of the alignment viewer allows sequences to be ordered according to the current tree layout while colour-coding nucleotide columns, enabling direct visual identification of matching and mismatching regions within the alignment. When the data are noisy, numerous small subtree rearrangements may occur and can be tedious to follow individually. The distance plot highlights windows with larger topological changes, helping to identify regions of interest. Across the full tree sequence, 1,403 subtree-move events were recorded. Summarising how often each leaf was part of a moved (highlighted) subtree, the largest contributions came from genotypes GII.4[P4], GII.4[P31], GII.4[P16], GII.17[P17], GII.2[P16], and GII.6[P7].

The complete SPR event frequency table, including per-taxon highlight counts, is provided with the example datasets and is publicly available in the project’s GitHub repository. Summarising our analysis shows that Phylo-Movies enables direct observation of recombination-associated changes in tree topology: as the sliding window crosses the ORF1/ORF2 junction, polymerase-defined clusters dissolve and capsid-defined clusters assemble. The alignment viewer further supports interpretation by reordering sequences according to the current tree layout and colour-coding nucleotide columns, allowing clustering patterns in the tree to be inspected alongside matching and mismatching regions in the alignment.

### 3.2 Visual Identification of Rogue Taxa in Bootstrap Tree Sets

Rogue taxa are leaves that cannot be reliably placed due to conflicting or missing data, causing them to shift position frequently across a set of reconstructed trees (Wilkinson, 1996). Their presence reduces resolution and support in consensus trees; removing them often yields more informative results. Identifying rogue taxa before final analysis is essential.

When analysing bootstrap trees, rogue taxa are those that change position across replicates, indicating unstable phylogenetic placement. Because Phylo-Movies visualises and highlights such moves, it provides a quick way to spot candidate rogues. To demonstrate this, we used two multiple sequence alignments from the RogueNaRok dataset (Aberer and Stamatakis, 2013b,a): Dataset 24 (24 taxa, 14,190 sites), corresponding to the palaeognath mitochondrial alignment analysed by Phillips et al. (2010), in which *Ostrich* has been identified as a rogue taxon due to unstable placement across bootstrap replicates; and Dataset 125 (125 taxa, 29,149 sites), whose taxa are anonymised (Seq1, Seq2, Seq3, etc.) and were contributed for benchmarking without disclosure of species identities, which we analysed to evaluate performance on a dataset larger than Dataset 24.

To order the resulting trees in a smooth trajectory that facilitates visual tracking of positional changes, we sorted replicates by their similarity in nucleotide composition to the original (non-resampled) alignment. For each replicate *b*, we computed the nucleotide composition **c**_*b*_ = (*n*_*A*_, *n*_*C*_, *n*_*G*_, *n*_*T*_, *n*_gap_) and measured its deviation from the original alignment using the Euclidean distance:

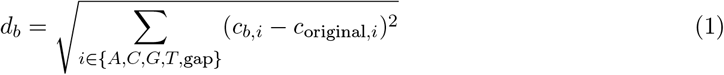

where *c*_original,*i*_ denotes the composition of the original multiple sequence alignment. Bootstrap replicates have no intrinsic biological order; any linear arrangement imposes a trajectory for visualisation. This does not alter the inferred trees themselves, but determines which pairs of trees are compared consecutively for the animation. The maximum-likelihood tree was then inferred from each bootstrap multiple sequence alignment using FastTree (Price *et al*., 2010) with GTR+Γ, employing the options `-gtr -gamma -nt -mlacc 2 -slownni -nosupport -quiet. The -mlacc 2` and `-slownni` settings increase optimisation accuracy by refining NNI moves and disabling heuristic subtree skipping, while SH-like support values were not computed.

We asked whether Phylo-Movies, when animating this trajectory of bootstrap trees, would highlight the same leaves identified as rogue taxa by RogueNaRok (Aberer and Stamatakis, 2013b). RogueNaRok identifies rogue taxa by optimising the relative bipartition information criterion (RBIC), which measures the overall support across a bootstrap tree set relative to the maximum achievable support. In contrast, Phylo-Movies does not compute a support-based criterion; instead, it decomposes differences between trees into sequential SPR moves and highlights the subtrees that change position across replicates.

In the 24-taxon dataset, based on 200 bootstrap trees, RogueNaRok reported an increase in RBIC from 0.828810 to 0.853333 upon removal of *Ostrich*. Correspondingly, *Ostrich* was the most frequently highlighted leaf in our SPR event frequency analysis, ranking 1st out of 24 subtrees (116 events; 18.33% of all highlighted splits; 95.8% percentile of SPR event frequency). The complete tables are available in our GitHub repository and can also be accessed directly through the example datasets provided within Phylo-Movies.

In the 125-taxon dataset, also comprising 200 bootstrap replicates, the taxon identified as rogue by RogueNaRok (*Seq112*) ranked 3rd out of 47 highlighted subtrees in our analysis and fell within the 94th percentile of highlight frequency (97 events; 6.75% of all highlighted splits).

Together, these results show that Phylo-Movies provides a rapid visual approach for identifying candidate rogue taxa, consistent with support-based rogue detection methods.

## Discussion

We have shown two use cases where visualising tree sequences is useful to gain insights that would be difficult to obtain by manual inspection. We illustrated this approach by visually identifying rogue taxa that substantially destabilize phylogenetic reconstruction, as their shifting positions across trees obscure otherwise stable branching patterns, and by inspecting a pronounced recombination breakpoint that underlies the differences observed between Norovirus genotypes.

Besides these two use cases, Phylo-Movies visualisation can be used in analysing many use cases involving tree sequences. A common application is the study of continuous topological changes along genomic alignments caused by recombination, a topic of particular interest in virology. When performing population simulations with recombination using tools such as the tskit framework (Kelleher *et al*., 2016), Phylo-Movies can be used to visualise the resulting tree sequence and to observe how recombination events alter genealogical structure along the genome. Moreover, if researchers have performed analyses showing that in some region there are changes in the topology or recombination breakpoints, Phylo-Movies can be used to visualise which subtrees or taxa might have been moving to result in the observed changes. However, there are no restrictions on the origin of the underlying multiple sequence alignment (MSA) from which the trees were inferred (e.g. bacterial or eukaryotic taxa), nor on the process that generated the resulting tree sequence.

Phylo-Movies can also be used to investigate additional aspects of tree variation along the underlying genomic sequence, including changes in phylogenetic resolution, instability of root or outgroup placement, branch-length variation, and other topology shifts not necessarily caused by recombination. Phylo-Movies also accepts multifurcating trees, expanding its range of applications. This allows users to observe how phylogenetic resolution varies across genomic regions—for instance, identifying windows where certain subtrees collapse into multifurcations due to insufficient informative sites, which may guide the choice of window size or indicate regions of genuine evolutionary uncertainty.

Similarly, one can check whether an outgroup used for rooting the trees remains stable, or whether the root branch ends in multifurcations (indicating loss of phylogenetic signal) or the outgroup jumps to different positions (showing rogue taxon behaviour). Both outcomes—loss of resolution and positional instability—show that the outgroup does not consistently root the tree across all genomic regions. Finally, Phylo-Movies is also able to visualise the change of branch lengths along the genomes even if the tree topologies remain the same during the whole range. Such changes may be caused by selection effects, which are sometimes misinterpreted as recombination events in purely distance-based analyses (e.g. Worobey *et al*., 2002).

A different application is possible when comparing trees inferred under different evolutionary models. Model misspecification can lead to systematic biases in topology recovery (Kapli and Telford, 2020), and Phylo-Movies could help visualise which subtrees are affected when the underlying substitution model changes. Similarly, when developing or analysing tree search strategies, one can use the visualisation to study the search trajectory through tree space, identifying which subtrees remain stable early versus those that continue rearranging until the final iterations.

When analysing tree sequences with many taxa, numerous trees, and frequent rearrangements, the distance plot helps locate regions of topological change so that users need not watch the entire animation to find interesting regions. For particularly large or complex datasets, it may be necessary to work with a subset of trees to manage memory and processing demands. This can be achieved by selecting a contiguous range of trees or by thinning the tree sequence to include only every *k*-th tree. As the number of taxa, trees, and rearrangements increases, the number of intermediate transitions that must be interpolated also grows, reducing visual clarity and making individual subtree movements more difficult to track. Although only 16 trees were inferred in the Norovirus analysis, the numerous and complex rearrangements between consecutive windows required 5,686 interpolated intermediate trees for visualisation. In general, increasing numbers of taxa, trees, and topological changes lead to more intermediate transitions, as each rearrangement must be decomposed into sequential subtree movements. This scaling behaviour represents a practical limitation, as computational and memory requirements increase with dataset size. Larger tree sequences therefore require the generation and rendering of more intermediate states, which may reduce responsiveness and visual clarity in highly complex analyses.

Like FigTree (Rambaut, 2006) and DensiTree (Bouckaert, 2010), Phylo-Movies reads trees in standard Newick format. However, whereas FigTree displays static images of individual trees and DensiTree summarises a set of trees by overlaying them, Phylo-Movies animates the topological changes between consecutive trees. In addition, Phylo-Movies can display two trees side by side, linking corresponding taxa with connecting lines—similar to a tanglegram(Page, 2002)—so that users can track the origin and destination of moving subtrees during the animation. In addition to animating topological transitions, Phylo-Movies provides functionality for alignment inspection, sliding-window tree inference, metadata-based colouring, and video export. These features support exploratory analysis and facilitate the interpretation of topological changes within a tree sequence.

## Data availability

All sequence alignments, phylogenetic trees, metadata, and code used in this study are available in the Phylo-Movies repository at https://github.com/enesberksakalli/phylo-movies. Phylo-Movies is released under the MIT License and includes a standalone desktop application. For the Norovirus example, corresponding GenBank accession numbers are provided in the repository metadata; accession information for the other examples is provided there as well.

The underlying transformation engine and backend logic are implemented in the `BranchArchitect` framework, which computes and decomposes topological differences between trees into sequences of rooted subtree prune-and-regraft (SPR) operations. The `BranchArchitect` source code is available at https://github.com/EnesSakalliUniWien/BranchArchitect.

Demonstration videos illustrating the Norovirus analysis and the rogue-taxon examples are available at: Norovirus (https://vimeo.com/1162400544); rogue taxon example 1 (https://vimeo.com/1162561152); rogue taxon example 2 (https://vimeo.com/1162563101).

## Competing Interests

The authors declare no conflict of interest.

## Funding

This work was supported by institutional funds from the University of Vienna. No external funding was received for this study.

## Notes

https://github.com/enesBerkSakalli/phylo-movies

### Competing Interest Statement

The authors have declared no competing interest.

